# Polar pattern formation induced by contact following locomotion in a multicellular system

**DOI:** 10.1101/2019.12.29.890566

**Authors:** Masayuki Hayakawa, Tetsuya Hiraiwa, Yuko Wada, Hidekazu Kuwayama, Tatsuo Shibata

## Abstract

Biophysical mechanisms underlying collective cell migration of eukaryotic cells have been studied extensively in recent years. One paradigm that induces cells to correlate their motions is contact inhibition of locomotion, by which cells migrating away from the contact site. Here, we report that tail-following behavior at the contact site, termed contact following locomotion (CFL), can induce a non-trivial collective behavior in migrating cells. We show the emergence of a traveling band showing polar order in a mutant *Dictyostelium* cell that lacks chemotactic activity. The traveling band is dynamic in the sense that it continuously assembled at the front of the band and disassembled at the back. A mutant cell lacking cell adhesion molecule TgrB1 did not show both the traveling band formation and CFL. We thus conclude that CFL is the cell-cell interaction underlying the traveling band formation. We then develop an agent-based simulation with CFL, which shows the role of CFL in the formation of traveling band. We further show that the polar order phase consists of subpopulations that exhibit characteristic transversal motions with respect to the direction of band propagation. These findings describe a novel mechanism of collective cell migration involving cell–cell interactions capable of inducing traveling band with polar order.

## Introduction

The collective migration of eukaryotic cells plays crucial roles in processes such as wound healing, tumor progression, and morphogenesis, and has been the focus of extensive study(Haeger et al., 2015). The collective effects are typically associated with cell–cell interactions, such as long-range interaction mediated by secreted chemicals or short-range stable cohesive interaction mediated by adhesion molecules. However, the study of self-propelled particles has revealed that motile elements which lack such activities may nonetheless give rise to dynamic collective motion, such as a traveling band(Chaté et al., 2008; Ginelli et al., 2010; Ohta and Yamanaka, 2014; Solon et al., 2015), mediated by a relatively simple transient short-range interaction, such as alignment interaction(Marchetti et al., 2013; Vicsek et al., 1995; Vicsek and Zafeiris, 2012). The emergence of such collective motions of self-propelled particles has been observed in a wide variety of systems, ranging from animal flocks(Ballerini et al., 2008), bacteria swarms(Wioland et al., 2013; Zhang et al., 2010), and cell assemblies(Szabó et al., 2006) to biopolymers and molecular motors(Butt et al., 2010; Sumino et al., 2012; Weber and Semmrich, 2010). For some of these systems, the connection between a macroscopic collective behavior and the microscopic dynamics of its constituents has been established. For instance, in biopolymers and molecular motors, traveling band formation is induced by local physical interactions among constituent elements(Sumino et al., 2012; Suzuki et al., 2015; Weber and Semmrich, 2010). In the case of eukaryotic cells, however, there has been no report to link traveling band formation to short-range cell–cell interactions.

The social amoeba *Dictyostelium discoideum* is a model organism for the study of collective cell migration. The coordinated movement of cell population is achieved by individual chemotactic motion to the cAMP gradient, which is formed in a self-organized way. However, a mutant cell that lacks chemotactic activity to cAMP still exhibits an organized coordinated motion that is probably mediated by cell-cell contacts(Kuwayama, 2013). Here, we demonstrate that this coordinated motion is a spontaneous polar order formation which phase-separates with a disordered background. We further show that this polar order formation is attributable to the tail-following behavior among the migrating cells, which we call contact following locomotion (CFL). We find that the polar ordered phase caused by CFL has an internal structure. An agent-based model with CFL further reveals that this internal structure is characteristic of the CFL-induced polar order formation. Thus, we establish the link between the collective behavior and the cell-cell interactions. Our findings open new possibilities that the concept of self-propelled particles contributes to the understanding of a highly orchestrated biological event of migrating cells in multicellular systems.

## Results

### Traveling band formation of non-chemotactic *Dictyostelium* cells

In the present study, we investigated collective cellular motion in a mutant strain of *Dictyostelium discoideum*, known as “KI cell,” which lacks all chemotactic activity(Kuwayama, 2013; Kuwayama et al., 1993), and thus does not form a cell aggregate under starvation conditions. Wildtype *Dictyostelium discoideum* forms an aggregate as a result of chemotaxis mediated by a self-secreted extracellular chemoattractant. Under starvation conditions, KI cells spread on a non-nutrient agar plate show a segregation of cell density, which propagates as bands in around six hours(Kuwayama, 2013), when the cell density is within a particular range (1.0 × 10^5^ cells cm^−2^ to 4.0 × 10^5^ cells cm^−2^) (Supplementary Movie 1). Initially, the traveling bands propagate in random directions with high orientational persistence. When two bands collide, they appear to pass through each other, retaining their shapes (Fig. 1a left) (Kuwayama, 2013). However, over time, the propagation directions gradually become aligned, probably due to weak reorientation of propagation direction as an effect of collisions. Finally, the bands are arranged almost periodically in space with a spatial interval of about 1 mm (Fig. 1a right, b).

**Figure 1.**
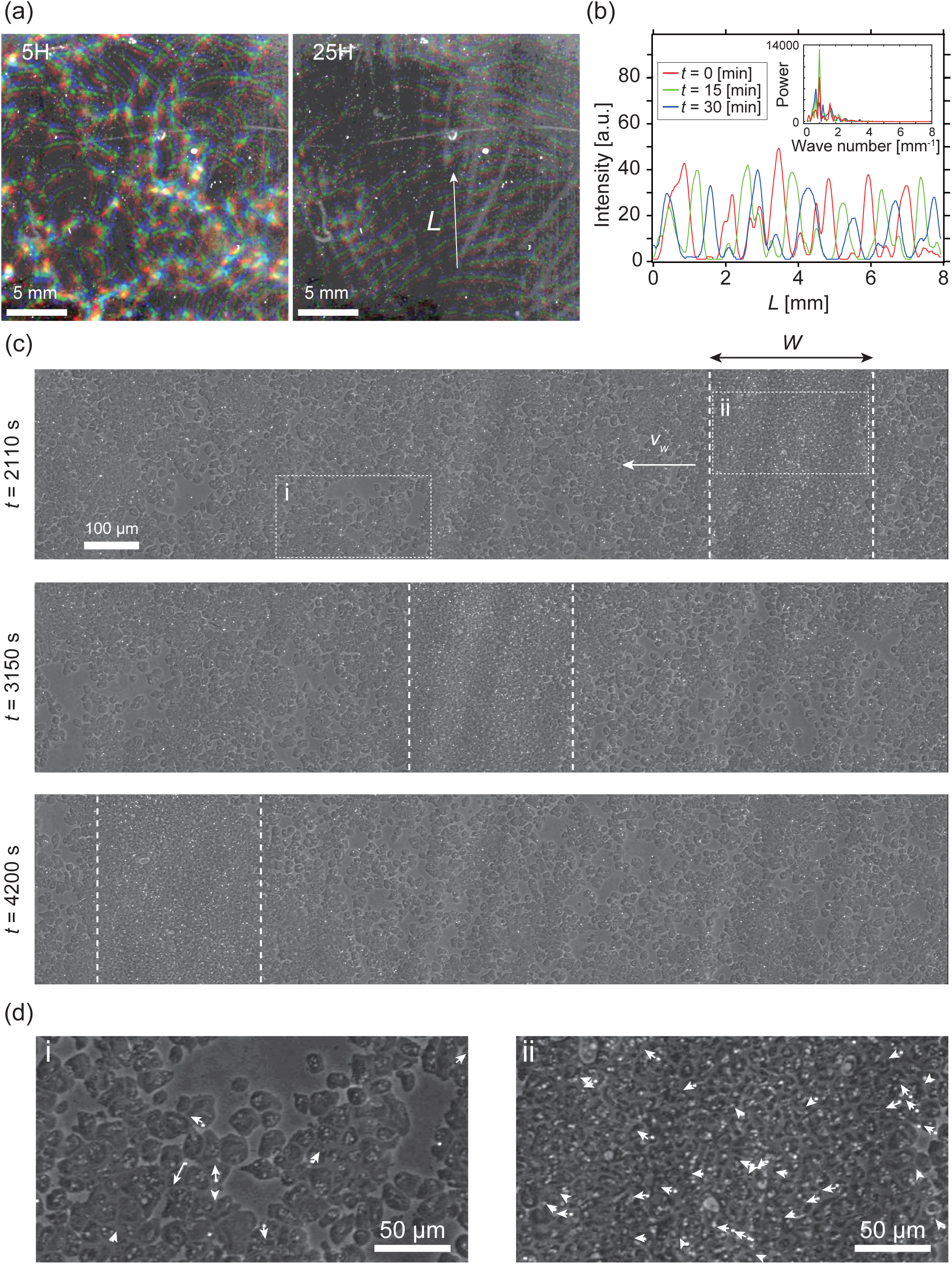
Segregation of cell density and formation of bands in non-chemotactic *D. discoideum* KI cell. **a**, The density profile of three time points with a time interval of 15 min indicated by color-coding (red *t*=0, green 15min, blue 30min). Brighter color indicates higher density. Five (left) and 25 (right) hours after incubation. See also Supplementary Movie 1. **b**, The intensity profile along the line indicated in (a), showing a periodic distribution of high-density regions. The inset shows a power spectrum of the intensity profile, indicating that the spatial interval was about 1 mm. **c**, Time evolution of phase-contrast image of high-density region (dotted lines) at *t*=2110, 3150, 4200 sec., respectively. The time points correspond to that in Supplemental Movie 2. **d**, High magnification images of low-density region (i) and high-density region (ii). Arrows indicate the migration directions of cells.

To determine the mechanism underlying this collective cellular motion, we conducted high-magnification observations. At around 16 hours after cells were seeded on an agar plate, a punched-out section of the agar plate was placed upside down on the glass slide, such that the monolayer of cells was sandwiched between agar and glass (Fig. S1a). These cells formed a high-density area that moved as a band in low-density area for long periods of time with high orientational persistence (Fig. 1c,d and Supplementary Movie 2). Whereas the cells in high-density area are packed without extra space, and thus the cell density is similar across different samples (Fig. S1c), the size *W* of the band along the propagation direction showed a broad distribution, ranging from *W =* 200 µm to 700 µm (N=10). In contrast, the traveling speed ⟨*ν*_*b*_⟩ *=* 0.5 ± 0.03 µm/s (N=10) was consistent among different bands, independent of size *W* (Fig. S1b).

### Analysis of single cell trajectories

To study the relationship between these collective behaviors and the migration of individual cells, we next performed cell-tracking analysis. Cellular movements were recorded by tracing the motion of fluorescent microbeads that were incorporated into the cells by phagocytosis. Figure 2a shows typical trajectories of individual KI cells. The distribution of migration speeds indicates that cell migration speed inside the band is slightly faster than that outside the band (Fig. 2b). The average migration speeds of individual cells inside and outside the band were *ν*_*in*_ *=* 0.38 ± 0.14 µm s^−1^ and *ν*_*out*_ *=* 0.30 ± 0.16 µm s^−1^, respectively. The migration direction of the cells inside the band was distributed around the direction of band propagation, while the migration direction outside the band was distributed almost uniformly (Fig. 2c). The mean squared displacement (MSD) inside the band was proportional to *t*^2^ for more than 10^3^ s (Fig. 2d). In contrast, the MSD outside the band exhibited a transition at around 100 s. from a persistent motion proportional to *t*^2^, to a random motion proportional to *t*, which indicates that this motion can be described as a persistent random motion with no preferred direction (Fig. 2d). This observed directional randomness reflects the effects of cellular collisions, as well as its intrinsic nature of single cells. In sum, cells inside the band exhibit directionally persistent motions, whereas cells outside move randomly.

**Figure 2.**
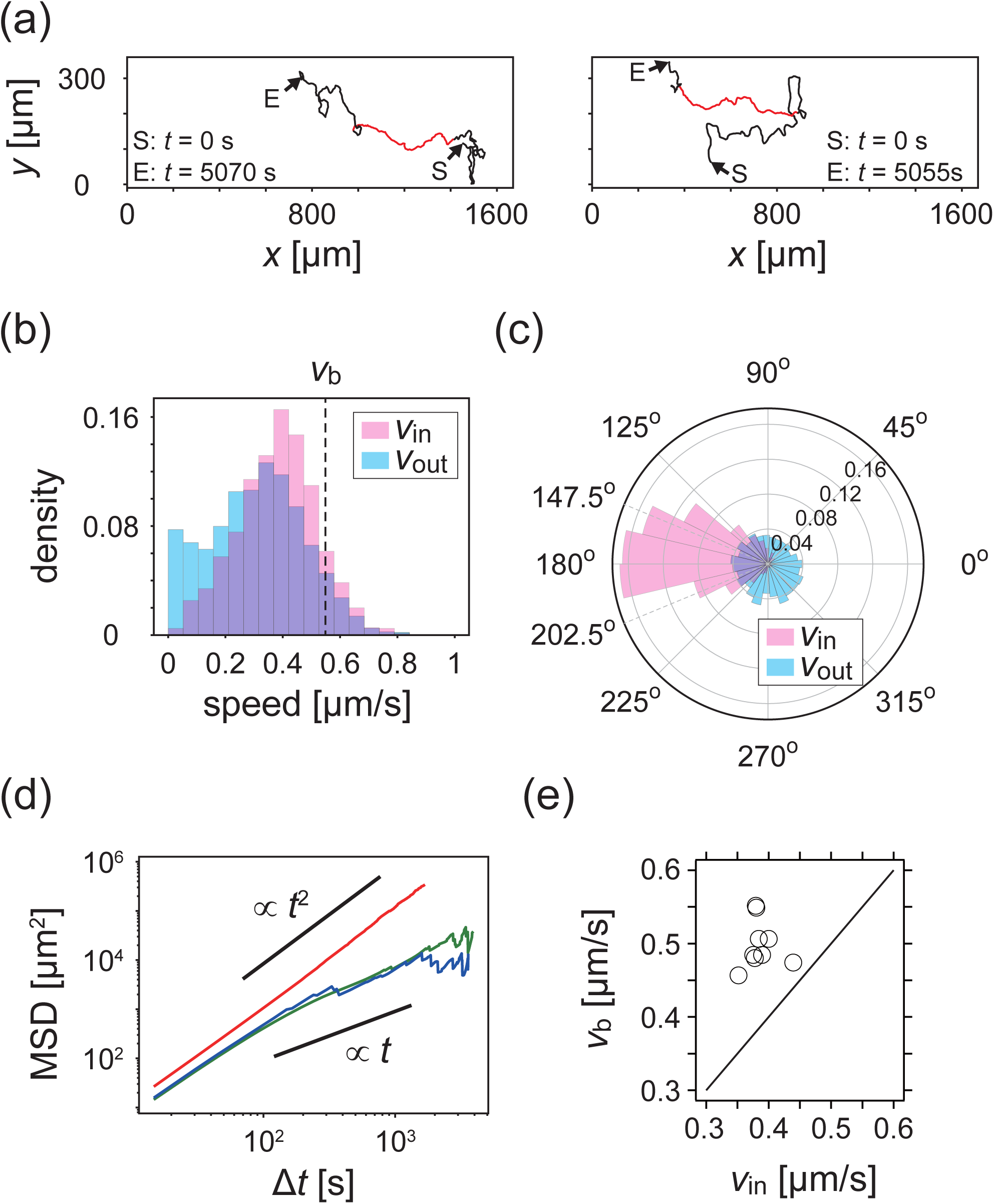
Analysis of single cell migrations inside and outside the band. **a**, Trajectories of single cells inside (red) and outside (black) the band. These trajectories were taken from the data shown in Fig. 1c. **b,c**, The distributions of the migration speed (b) and migration direction (c) inside (pink) and outside (blue) the band. **d**, Mean squared displacement (MSD) of cell motions inside the band (red), before entering the band (green) and after leaving the band (blue) **e**, Scatter plot of the band speed *v*_*w*_ against the cell speed ⟨*v*_*in*_⟩ within the band. The number of bands investigated is *N* = 10.

### Propagation of cell density profile

We then compared the average cell speed inside the band *ν*_*in*_ and the band propagation speed *ν*_*b*_, (Fig. 2e), and found that the band propagates faster than the cell migration speed for all samples investigated. This implies turnover of cells in the band, and that the band is continuously assembled at the front of the band and disassembled at the back. Thus, it is the cell density profile that shows propagation as a band(Kuwayama, 2013).

### Analysis of multicellular movement reveals polar order formation

To quantitatively characterize the multicellular movement, we introduce the local polar order parameter, *φ*(*n, t*) *=* |⟨***ν***_*i*_ (*t*)/|***ν***_*i*_ (*t*)|⟩_*i*∈ℒ(*n*)_ |, obtained from the instantaneous cell velocity ***ν***_*i*_ (*t*), where ℒ(*n*) is the *n*th domain along the direction of band propagation (see Supplementary text). In the high-density region that propagates as a band, *φ*(*n, t*) reaches around 0.8, while *φ*(*n, t*) in the low-density area remained below 0.4 (Fig. 3a). Thus, the high-density region is polar-ordered phase, which propagates in the low-density disordered phase. The polar order parameter of the band showed intersample variability, and was distributed from 0.6 to 0.85 (Fig. 3b). We found that the order parameter of band was positively correlated with the width of band *W* (Fig. 3b).

**Figure 3.**
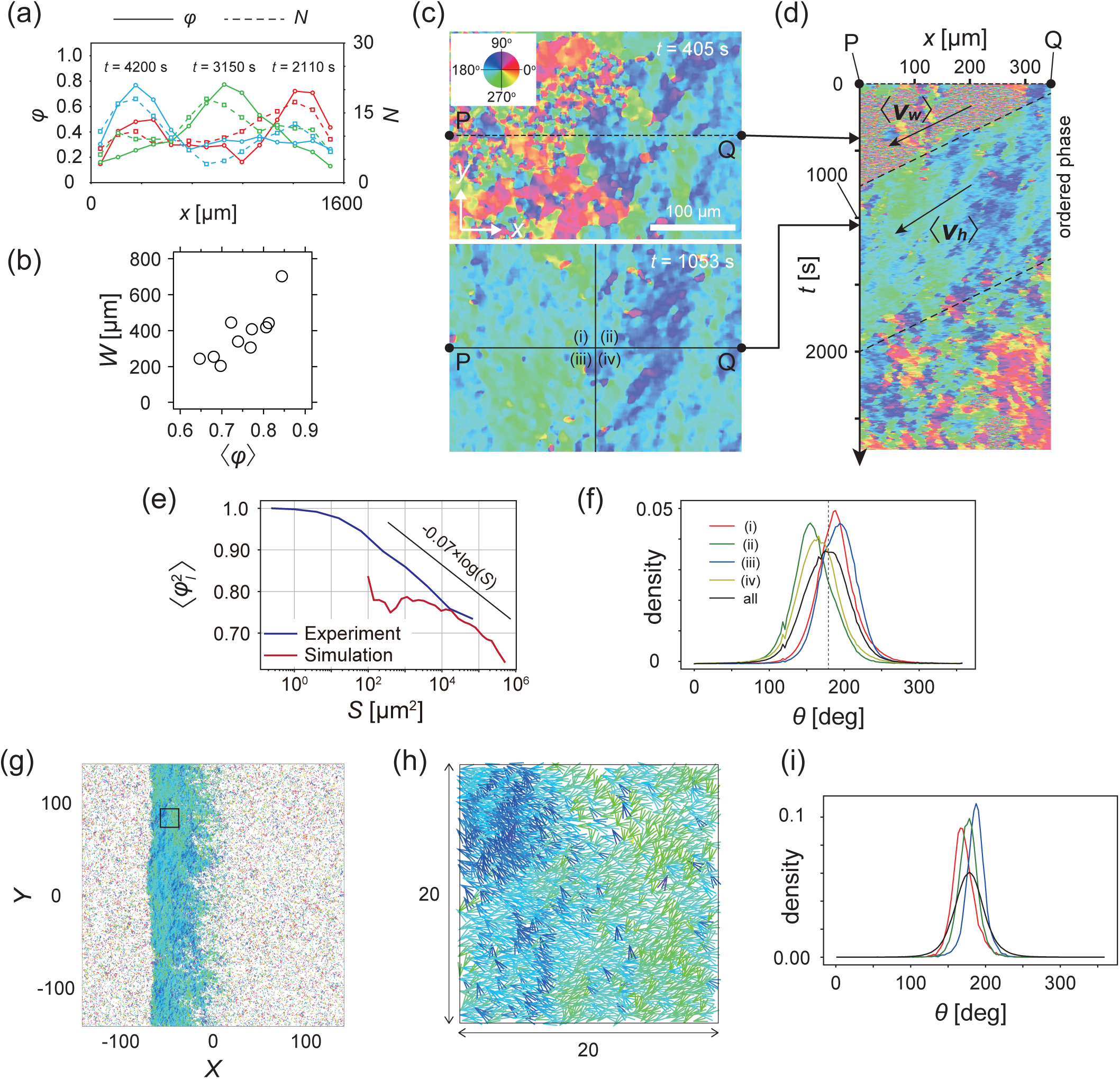
Analysis of heterogeneity within the ordered phase. **a**, Spatial profile of the local polar order parameter (solid lines) and the number of beads in the intervals (dotted lines) in Fig. 1c. **b**, Scatter plot of band width against polar order parameter within the band region. The number of bands studied is *N* = 10. **c**, Optical flow images in the front region of a band *t*=405 (top) and within the band *t*=1083 (bottom). See also Supplementary movies 3 **e**, Kymograph of the optical flow image along the line PQ shown in c (top). The arrows indicate the average velocity of band ⟨*v*_*b*_⟩ and the average cell speed ⟨*v*_*in*_⟩. **e**, Size-dependent squared local order parameter plotted against the area ***S*** for the data shown in **c. f**, Probability distribution function (pdf) of the migration direction within the band region obtained by the optical flow analysis shown in c. The pdfs (i)-(iv) are obtained in the regions (i)-(iv) in c (bottom), respectively. Average pdf is shown by the black line. **g**, Snapshot of simulation result showing a polar ordered phase as a propagating band in the background of disordered phase. The color code indicates the migration direction of individual particle as shown in **c**. See also Supplementary movies 8. **h**, Magnification of squared area shown in **g**. The size of area (20×20) is comparable to the whole area shown in **c**. Each arrow indicates the direction of polarity. **i**, Probability distribution function of the migration direction within the band region in the simulation (red, green and blue lines). For the choice of ROI, see Supplementary text. Average pdf is shown by the black line.

### Internal structure in the polar ordered region

The polar order phase is not completely homogeneous with respect to migration direction, but exhibits heterogeneity; this is related to the underlying assembly mechanism. This heterogeneity can be visualized in the velocity field obtained by optical flow, in which the direction of cell migration can be distinguished by color (Fig. 3c and Supplementary movie 3). The size-dependent squared local order parameter 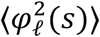 (see Supplementary text) shows a logarithmic decay with area *S* (Fig. 3e), indicating that this heterogeneity is not spatially uncorrelated. Within the band (Fig. 3c bottom), the migration direction was widely distributed from about 145 to 210 degrees (a black line in Fig. 3f). The probability density functions (pdf) of the migration direction obtained for the four regions (Fig. 3c bottom (i–iv)) show peaks at different directions (Fig. 3f), indicating the presence of two subpopulations; one in which the migration direction is ∼160° (regions (ii) and (iv)) and another in which it is ∼190° (regions (i) and (iii)). These two subpopulations are also recognized in Fig. 3c (bottom) as the regions with dark blue and light green colors, respectively, forming stripes. These two types of stripes extend perpendicular to the direction of band propagation, and are alternately arranged. The typical width of the stripe was around 125 µm, as determined by the analysis of autocorrelation function (Fig.S2a). The kymograph in Fig. 3d shows the temporal evolution of the velocity field along the line PQ in Fig. 3c, indicating that the stripes (light green and dark blue) are almost immobile, suggesting that the same cells experience the two stripes sequentially. From a reference frame co-moving with the band, cells move from the front to the end, during which they move downward in a stripe, and then move upward in another stripe (Fig.S2b). These analyses illustrate that the polar order phase possesses an internal structure with respect to the migration direction.

### Contact following locomotion is the cell-cell interaction that induces polar pattern formation

The formation of a polar-ordered phase with an internal structure is ultimately related to the microscopic interactions between individual cells, which are short-ranged. In the low-density region, cells are not completely isolated, but rather are often associated with each other, migrating in single files (Fig.4a and Supplementary movie 4). This tail-following behavior has been described for wild-type *Dictyostelium* cells within aggregation streams(Dormann and Parent, 2002). We call this behavior “contact following locomotion” (CFL). In low-density assay, when two cells collide, they either form CFL (Fig. 4b and Supplementary movie 5) or not (Fig. S3a). To quantitatively characterize CFL, we measured the duration of cell–cell contact after two cells collide. During the formation of CFL, the typical cell-to-cell distance is given by *d*_*a*_ *=* 24 µm. We measured the time interval during which the distance is less than *d*_*a*_ from the time series of the distance between two cells (Fig. S3b). As shown in Fig.4c, in half of the cases, cell–cell contact persists for more than 300 sec. To determine whether cells that form contacts for >300 sec exhibit tail-following behavior, we measured the average angle *a* of the angles *a*_1_ and *a*_2_, which are the angles of the velocity vectors ***ν***_1_ and ***ν***_2_ with respect to the vector connecting the two cell centers ***d***, respectively (Fig. 4d). In almost 60% of all cases, the angle *a* is 0–30° (Fig. 4e), indicating tail-following type CFL.

**Figure 4.**
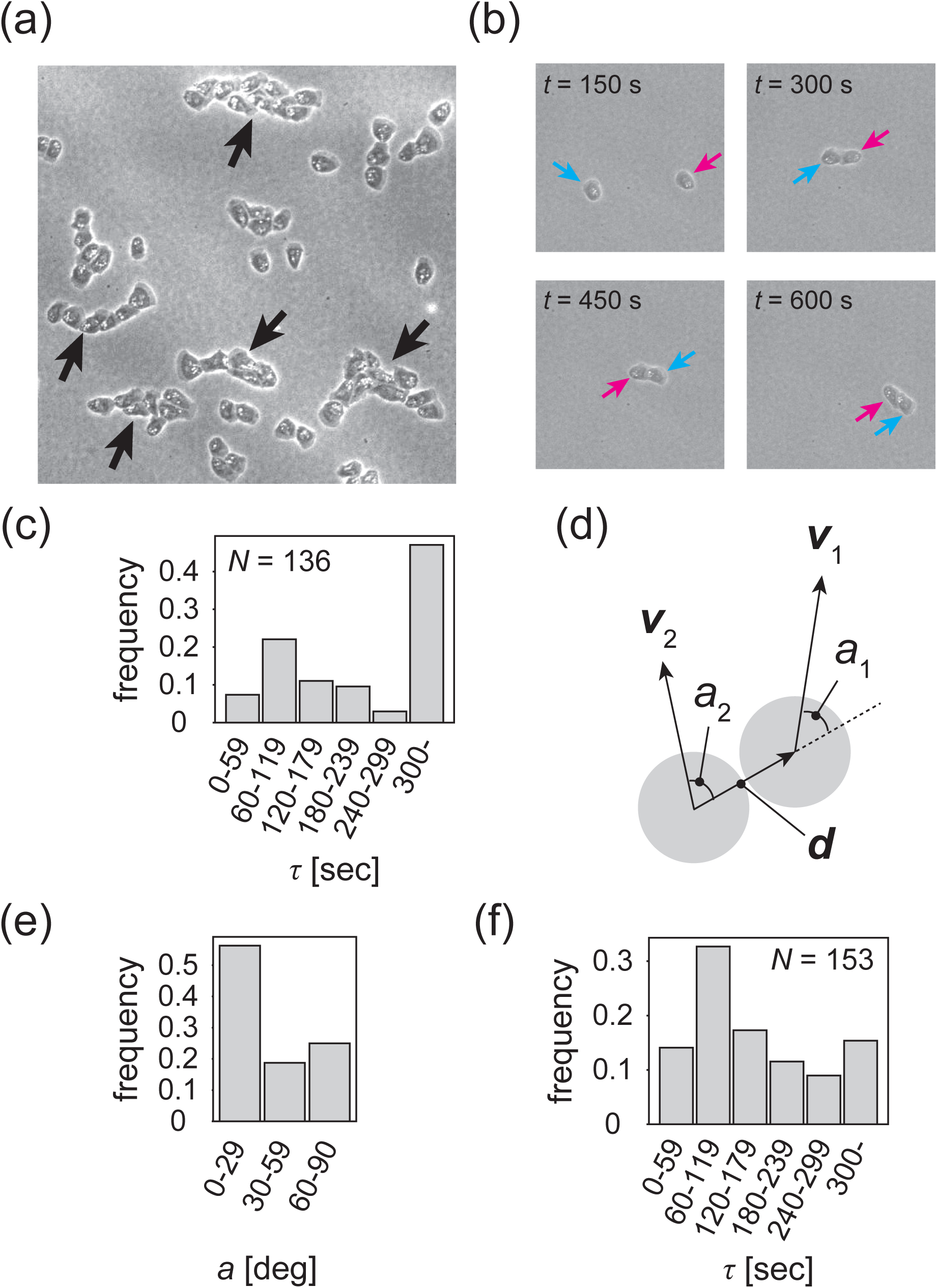
Contact following locomotion responsible for band propagation. **a**, Snapshot of the contact following locomotion. See also Supplementary movies 4. **b**, Representative time evolution of collision of two cells. Colored arrows represent the same cell. See also Supplementary movies 5. **c**, Histogram of the duration of two cell contacts for KI cell (control). **d**, Schematic of angle *a*_1_ (*a*_2_), which is the angle of the velocity vector ***v***_1_ (***v***_2_) with respect to the vector ***d*** connecting two cell centers. Then, the angle *a* is obtained as the angular average of *a*_1_ and *a*_2_, i.e., *A* cos *a* = (cos *a*_1_ + cos *a*_2_)/2, *A* sin *a* = (sin *a*_1_ + sin *a*_2_)/2. **e**, Histogram of the angle *a* for the KI cells that contact each other for >300 sec. **f.** Histogram of the duration of two cell contacts for the *tgrb1* null mutant.

To determine whether CFL is responsible for the collective behavior of KI cells, we sought a mutant cell that lacks CFL activity. A knockout mutant that fails to express the cell–cell adhesion molecule TgrB1 exhibits reduced CFL activity(Fujimori et al., 2019). TgrB1 is known to mediate cell–cell adhesion via a heterophilic interaction with its partner TgrC1(Fujimori et al., 2019; Hirose et al., 2011; 2015; C.-L. F. Li et al., 2015). We first assessed whether the *tgrb1* null mutant forms propagating bands. As in the control case, under starvation conditions, we spread the *tgrb1* null mutant cells on a non-nutrient agar plate at a cell density of 2.0 to 3.0 × 10^5^ cells cm^−2^ (see Methods). However, neither segregation of cell density nor propagating bands appeared (Supplementary movies 6 and 7.).

We then compared locomotive activity between control cells and *tgrb1* null mutants. The velocity auto-correlation functions *C*(*τ*) of the isolated single cells showed similar behaviors (Fig. S3c), indicating that locomotive activity was comparable between KI cells and the *tgrb1* null mutant cells. We next quantitatively characterized the formation of cell–cell contacts. We found that in 80% of all cases, cell–cell contact is disrupted before 300 sec (Fig. 4f), and that only 10% of cells established CFL (Fig. S3d). In particular, in half of all cases, the cell–cell distance becomes larger than *d*_*a*_ in 120 sec, indicating that these cells failed to establish cell–cell contact. Our analyses illustrate that in the *tgrb1* null mutant, CFL is nearly absent. We conclude that CFL is essential for the segregation of cell density and the formation of propagating bands.

### Mathematical modeling of polar pattern formation driven by contact following locomotion

The collective motion of KI cells induced by the CFL interaction can be modeled by an agent-based simulation(Hiraiwa, 2019). In the model, particle *i* at position **r**_*i*_ self-propels at a constant velocity *ν*_0_ in the direction of its own polarity **q**_*i*_ subjected to white Gaussian noise. Thus, without interactions, the particles exhibit a persistent random walk(Hiraiwa et al., 2014). The effect of CFL is introduced so that polarity **q**_*i*_ orients to the location of the adjacent particle *j*, when particle *i* is located at the tail of particle *j* (parameterized by *ζ*). In addition to this effect, the particles interact with each other through volume exclusion interaction, adhesion, and the effect of polarity **q**_*i*_ orienting toward the direction of its velocity **v**_*i*_ *= d***r**_*i*_/*dt* (parameterized by *α*). For a fixed parameter set (*α =* 0.4; see Supplementary text), without CFL (*ζ =* 0), the collective behaviors did not form (Fig. S4). In contrast, with CFL (*ζ* ≥ 0.1), a polar-ordered phase appeared as a propagating band in the background of disordered phase (Fig. 3g and Supplementary movies 8). The speeds of the traveling band and particles within the band were 0.96 and 0.9, respectively, relative to the speed of isolated particles, indicating that the band is dynamic with assembly in the front and disassembly in the tail, consistent with our experimental results. From the spatial pattern shown in Fig. 3gh, in which the migration direction is indicated by color code, heterogeneity in the migration direction is recognized within the polar-ordered phase. In the simulation, we studied the pdf of migration direction in regions, whose size is comparable to that in Fig. 3c ((i)–(iv)), and found that the pdf exhibited peaks at different directions (Fig. 3i), similar to our experimental results (Fig. 3f). To determine whether this formation of internal structure is a characteristic of propagating bands induced by CFL, we studied a propagating band formed by increasing alignment effect *α* without CFL (*ζ =* 0), and found that the pdfs of migration direction exhibit peaks at closely similar positions, indicating that the migration direction in the ordered phase is more homogeneous (Fig. S5e). Thus, the formation of internal structure appears to be a characteristic of the collective behavior induced by CFL. The size-dependent squared local order parameter 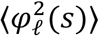 (see Supplementary text) also shows the characteristic decay with a logarithmic dependence on area *S* (Fig. 3e), as observed experimentally.

## Discussion

In this study, we report that a mutant of *Dictyostelium* cell that lacks all chemotactic activity exhibits spontaneous segregation into polar ordered solitary band. This pattern formation is attributable to the cell-cell interaction called contact following locomotion (CFL). The agent-based model that includes CFL reproduces the observed macroscopic behaviors. Thus, we establish a link between the microscopic cell-cell interactions and the macroscopic polar pattern formation.

We showed that the width of band is distributed widely from *W =* 200 µm to 700 µm (Fig. S1b). The local cell density within the band is similar across different samples (Fig. S1c), suggesting that the local cell density may not be a relevant factor. In contrast, we found the positive correlation between the width and the order parameter within the band (Fig. 3b). We speculate that if the correlation in the migration direction is gradually decorrelated from the front to the end of the band, bands with lower order parameters will be more prone to larger decorrelation in the migration direction. Consequently, we expect that the stronger the polar order, the wider the band width *W*.

One characteristic behavior of the present polar pattern formation is the formation of internal structure, which consists of subpopulations with transversal motions. From the numerical simulation result, this formation of subpopulation was not seen in the model without CFL. Thus, the internal structure is a characteristic of CFL induced polar pattern formation. Resolving how fluctuations in the migration direction perpendicular to the band propagation direction grow to form subpopulations when CFL is present remains a question for future study.

In this paper, we mainly focused on the behavior of single solitary band. We studied the traveling band, which was well separated from other bands. Thus, all properties of single solitary band studied in this paper is independent of interaction between different bands. In some area, the traveling bands are arranged almost periodically in space with a spatial interval of about 1 mm (Fig. 1b). How bands interact with each other to reach a periodic spacing and whether the interval is independent of band width *W* are to be investigated.

Wildtype *Dictyostelium discoideum* usually aggregates through chemotaxis to form a hemispherical mound with a central tip region that regulates the formation of slug-like multicellular structure(Williams, 2010). Whereas the KI cell alone does not show this activity, KI cells are able to spontaneously migrate to the central tip region transplanted from a wildtype slug and undergo normal morphogenesis and cell differentiation; this is not observed in mutant KI cells lacking tgrB1(Kida et al., 2019), suggesting that tgrB1-dependent CFL without chemotaxis allows KI cells to spontaneously migrate in slug. Furthermore, in wildtype cells, the chemical guidance cue has been shown to cease during the multicellular phase, which suggests that an alternative mechanism induces collective cell migration in the multicellular body(Hashimura et al., 2019). We propose that polar order formation induced by CFL plays an important role in late-stage morphogenesis in this organism. Contact following locomotion, or chain migration, have been reported in other cell types(D. Li and Wang, 2018). The macroscopic behaviors reported in this paper may thus be found in other systems as well.

## Materials and Methods

### Culture condition of KI mutant cells and cell density measurement

1 mL of *Klebsiella aerogenes* suspended in 5LP medium (0.5% Lactose, 0.5% bactopeptone 211677, Optical density = 0.1) was spread on the 9 cm 5LP plate (0.5% Lactose, 0.5% bactopeptone 211677, 1.5% agar), 5LP medium dried, the non-chemotactic *Dictyostelium discoideum*, KI mutant cells were inoculated on the plate. The KI cells were incubated for about five days at 21 °C. After cultivation, the KI cells and *Klebsiella* on the plate were collected with a phosphate buffer (PB). To remove the *Klebsiella*, the suspension was centrifuged and discard as much of the supernatant liquid as possible by aspiration, then clean PB was added. After repeating this process two times, the number of cells was counted using a hemacytometer.

### Macroscopic observation of the traveling bands

The washed KI cells were spread (cell density = 5.0×10^5^ cells/cm^2^) on a 9 cm non-nutrient agar plate (1.5% agar) to cause starvation. After drying of the PB, the plate was scanned every 15 minutes using a film scanner (V850, EPSON). The brightness in scanner images is inversely correlated with cell density(Takeuchi et al., 2014). For Fig 1a and b, the original images were inverted with color that depends on time points.

### Microscopic observation of the traveling bands

The KI cells were spread (cell density = 2.0 to 3.0×10^5^ cells/cm^2^) on the non-nutrient agar plate and incubated at 21 °C for around 16 hours. A punched piece of the agar plate was placed upside down on the glass slide, and the travelling bands between the agar and glass was observed by phase contrast imaging using an inverted microscope (TiE, Nikon with a 20x phase-contrast objective, equipped with an EMCCD camera (iXon+, Andor)).

### Tracking analysis of individual KI cells

For the tracking analysis, 1 μL of the PB including 3% fluorescent microbeads (ex:441, em:486, 1.0 μm, Polysciences, Inc.) was spread at the same time with the KI cells. The trajectories of the microbeads were automatically tracked by using the ParticleTracker 2D, a plugin for Image J (National Institutes of Health, USA). To eliminate the trajectories of the microbeads that was not internalized by the KI cells, if |***ν***_cell_| was slower than 0.25 μm/s for 300 second continuously, we excluded such trajectories. We also excluded the short trajectories of which continuous tracked time was shorter than 1 hour.

### The mean squared displacement (MSD)

The MSD (Fig.2d) was calculated using the formula below.

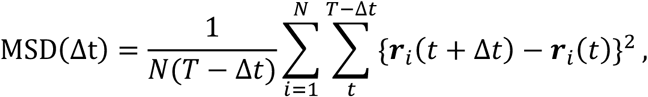

where *Δt, T*, and *N* means a time interval, final time, and number of the trajectory, respectively.

### Local polar order parameter

To obtain the local polar order parameter *φ*(*n, t*) shown in Fig. 3a, the picture shown in fig 1c was divided into *n* sections with width *Δx* (µm), and the order parameter was calculated in each section at each time from the trajectories obtained by the tracking analysis. The local order parameter *φ* is defined as

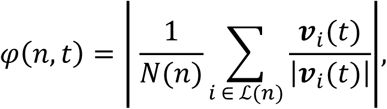

where ℒ(*n*) is the set of cells that satisfy (*n* − 1)*Δx* ≤ *x*_*i*_ ≤ *nΔx, N*(*n*) is number of the cells in ℒ(*n*), ***v***_i_ and *x*_*i*_ are the velocity and *x*-position of *i*-th fluorescent microbeads, respectively. In this study, *n* = 14 and *Δx* = 119 µm.

### Optical flow analysis

Optical flow analysis was performed based on the Gunnar-Farneback method using OpenCV library. In the optical flow analysis, the displacement of each pixel in the original pictures are characterized by coloring based on the HSV (“hue”, “saturation”, “value”) representation. The “hue” varies depending on angular variation of each pixel. In this study, a “saturation” and “value” of the processed images via optical flow was fixed to 150 and 255, respectively. The sequential images of the traveling band used for this analysis were taken every 2 seconds.

### Size-dependent squared local order parameter

To characterize the internal structures of the traveling band, the size-dependent squared local order parameter 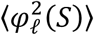 is introduced (Fig.3e). To obtain the size-dependent squared local order parameter, we first calculate the squared polar order parameter 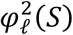 within a ROI of size *S*, which is defined as

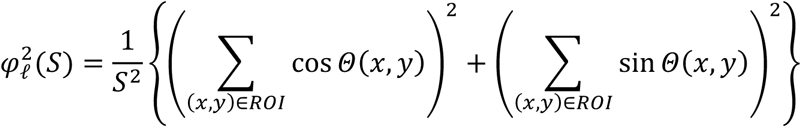

where *Θ*(*x, y*) = *hue* × (360/255) indicates the angular variation of the pixel at position (*x, y*). The value of *hue* was obtained from the optical flow analysis (Fig. 3c). Then, 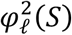 is averaged over the entire area to obtain 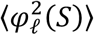. If *Θ*(*x, y*) is a random number without spatial correlation, as the increase of the area *S*, 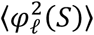 is expected to decay in proportion to *S*^−1^.

### Autocorrelation function of transverse motion with respect to the band propagation direction

Because the band show propagation in *x*-direction, autocorrelation function of transverse motion *C*_sin_ is defined using *y*-component of motion as

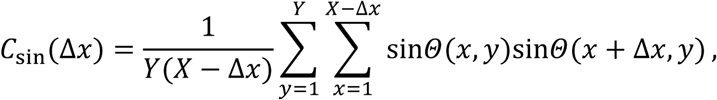

where Δ*x* is pixel interval along the *x*-axis. *C*_sin_ was plotted after that unit of Δ*x* is converted to the length.

### Preparation of *tgrb1* null mutant cells

The gene disruption construct for *tgrB1* was synthesized by a polymerase chain reaction (PCR)-dependent technique (Kuwayama et al., 2002). Briefly, the 5-flanking region of the construct was amplified with two primers, 5-CAACAGGTGGAGACTTCGGG-3 and 5-GTAATCATGGTCATAGCTGTTTCCTGCAGGCCAGCAGTAATAGTTGGAG-3. The 3-flanking region of the construct was amplified with primers, 5-CACTGGCCGTCGTTTTACAACGTCGACGAGAACTGTTGATTCTGATGG-3 and 5-CTTGGTCCTGAACGAACTCC-3. The bsr cassette in the multicloning site of pUCBsr Bam [Adachi et al., 1994] was amplified using the primer pair 5-CTGCAGGAAACAGCTATGACCATGATTAC-3 and 5-GTCGACGTTGTAAAACGACGGCCAGTG-3, both of which are complementary to the two underlined regions, respectively. The three amplified fragments were subjected to fusion PCR that produced the required gene-targeting construct. The gene-targeting constructs were cloned using a TOPO TA cloning kit for sequencing (ThermoFisher Scintific MA, USA).The linear construct was amplified by PCR using the outermost primers up to 10 µg. and transformed into KI-5 cells. The KO clones were selected by genomic PCR using the outermost primers.

### Culture condition and starvation treatment of *tgrb1* mutant null cells

The *tgrb1* null cells were cultured in HL5 medium (1.43% Proteose Peptone 211684, 0.72% Yeast Extract212750, 1.43% Gulcose, 0.05% KH_2_PO_4_, 0.13% Na_2_HPO_4_12H_2_O) at 21 degree Celsius. After reaching confluent, cells on the bottom were peeled off and collected, then washed two times with a centrifuge and PB. Next, the *tgrb1* null cells were transferred on the 1/3 SM plate (0.33% Gulcose, 0.33% bactopeptone 211677, 0.45% KH_2_PO_4_, 0.3% Na_2_HPO_4_, 1.5% agar) with *Klebsiella* suspension, and incubated for around two days at 21 °C. After, through the wash and count, the *tgrb1* null cells were spread on the non-nutrient agar plate, after which the plate was scanned every 15 minutes using the film scanner.

### Characterization of the contact following locomotion

The KI cells and *tgrb1* null cells for the collision assay were scraped from the traveling bands and surface of the plate, respectively. The scraped cells were placed on the non-nutrient agar and sandwiched with the glass. After around one hour incubation at 21 °C, binary collisions of two cells were observed by microscopy and recorded every 15 seconds. The motion of the cells was tracked manually using the Manual Tracking, a plugin of Image J. Here, collision was defined as the contact of pseudopods.

### The velocity autocorrelation function

Firstly, the migrations of the KI and *tgrb1* null mutant cells were recorded every 20 s for 60 min. Here, to extract an intrinsic locomotive activity of the cells, interactions with other cells, wall, and etc. were eliminated. Using obtained trajectories of cells that migrate with the velocity ***v***, the velocity autocorrelation function *C*(*τ*) was calculated. *C*(*τ*) is described with the form of

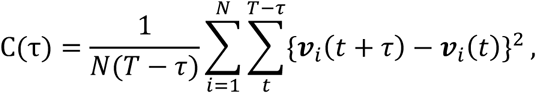

where *τ, t, T*, and *N* means a time interval, time, final time, and number of the trajectory, respectively (Fig. S3c).

### Modeling collective motion induced by contact following locomotion

The collective motion of KI cells induced by the CFL interaction can be modeled by an agent-based simulation. In the model, self-propelled particle *i* at position **r**_*i*_ moves at a constant velocity ν _0_ in the direction of its own polarity **q**_*i*_ subjected to white Gaussian noise. Thus, without interactions, the particles exhibit persistent random walk(Hiraiwa et al., 2014). Collective motion can be modeled by assuming particle-particle interactions(Hiraiwa, 2019). We firstly assume that the particles interact with each other through volume exclusion (parameterized by *β*) and adhesion (parameterized by *γ*). We also assume the feature that the polarity of each particle orients to the direction of its velocity **v**_*i*_ = *d***r**_*i*_/*dt* (parameterized by *α*); it is known that this assumption can effectively give rise to the alignment interaction between the particles when it is combined with the volume exclusion effect(B. Li and Sun, 2014). (Therefore, we simply refer to this feature as alignment effect in the main text.) As the main focus of this article, we incorporate CFL into this model by assuming the particle-particle interaction by which polarity **q**_*i*_ orients to the location of the adjacent particle *j* when particle *i* is located at the tail of particle *j* (parameterized by *ζ*). The equation of motion for the particle *i* are then given by

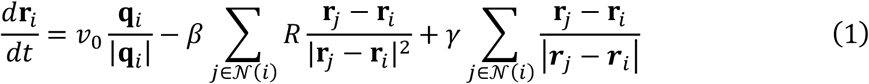

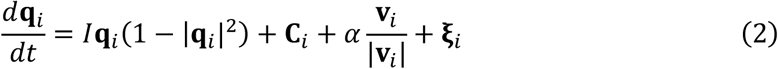

where the second and third terms on the right-hand side of Eq.(1) are the effects of volume exclusion and adhesions, respectively. Here, *𝒩*(*i*) is a set of particles that are contacting with the particle *i*, i.e., the particle *j* ∈ *𝒩*(*i*) satisfies |**r**_*j*_ − **r**_*i*_| ≤ *R*. On the right hand side of Eq.(2), the first term shows the self-polrization, the third term gives the effect that the polarity orients to the velocity direction **v**_*i*_/|**v**_*i*_|, the last term is white Gaussian noise with ⟨**ξ**_*i*_⟩ = (0,0) and ⟨**ξ**_*i*_(*t*). **ξ**_*j*_(*t*′)⟩ = *σ*^2^*δ*_*ij*_*δ*(*t* − *t*′), and the second term **C**_*i*_ describes the CFL, parameterized by *ζ*, given by

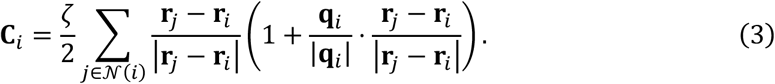

Here, when the polarity of particle *j*, **q**_*j*_, and the vector from particles *i* to *j*, **r**_*j*_ − **r**_*i*_, are in the same orientation, the maximum following effect is exerted on particle *i* to the direction of particle *j*. Such a situation is expected when particle *i* is located in the tail of particle *j* with respect to the polarity **q**_*j*_. In contrast, when particle *i* is located in the front of particle *j*, **C**_*i*_ almost vanishes. The simulation is implemented within a square box of size *L* with periodic boundary condition. For all simulations, we used fixed parameter values except *ζ* and *α*, given by *v*_0_ = 1.0, *β* = 1.0, *R* = 1.0, *γ* = 1.20, and *σ*^2^ = 0.4. For *I* in Eq.(2), we consider the situation where *I* is infinitely large, so that ***q*** was projected onto the unit vector |***q***| = 1 for the numerical simulation. The density of particles per unit area *ρ* is given *ρ* = 1. The number of particles *n* is *n* = 80,000 (Fig. 3ghi and Fig. S5) and *n* = 10,000 (Fig. S4).

### Histogram of migration direction in the numerical results

Firstly, we selected only the ROIs in the vicinity of the band front in the following way: We define a ROI as being within the bands if the particle density is higher than 1.16, which corresponds to the 2D dense packing fraction of disks, ∼0.91. Using this definition, we define the ROI as being vicinity of the band front if the ROI is within the band at the last *F*_*ana*_ frames whereas it is out of the band at the frames between the last *F*_*ana*_ + *F*_*wait*_+ *F*_*out*_ and the last *F*_*ana*_ + *F*_*wait*_. In other words, *F*_*ana*_ means the number of frames to be analyzed and must be within the band, *F*_*out*_ means the number of frames to determine the band front (i.e. the frames in which the ROI must be still out of the band assuming that the band travels only in one direction), and *F*_*wait*_ means the number of the waiting frames (i.e. the frames which are not used at all) between these frame sets.

The results of this algorithm for CFL-induced (*ζ* = 0.1, *α* = 0.4) and alignment-induced (*ζ* = 0.0, *α* = 1.0) bands are shown in Figs. S5a and b, respectively. Here, we used the following sets of the parameters for our analysis in this article: *F*_*ana*_ = 55, which corresponds to the time window in the analysis of experimental data. *F*_*wait*_ = 76, with around which the band front can propagate across one ROI. *F*_*out*_ = 5, which has been empirically determined. The duration between each frame is *dt* = 0.2 in the unit of time of our numerical simulation.

Secondly, using these near-front ROIs, we calculate the histograms of migration direction (d**r**_*i*_(t)/dt)/|d**r**_*i*_(t)/dt| for each ROI using all the *F*_*ana*_ frames (t) and the particles (*i*) in it at each frame. Then, we plot only the histograms for the ROIs which have the top eight and nine peak probability densities for CFL-induced and alignment-induced bands, respectively. The results are plotted in Figs. S5c and d, respectively. One can find the clear difference in these histograms between the CFL-induced and alignment-induced bands. The peak position and height for the CFL-induced band have large varieties, whereas those for alignment-induced band are less distributed. Furthermore, the peaks for the CFL-induced band are much higher than those for alignment-induced band. Figure 3i of the main text and Fig. S5e plot three typical histograms from Fig. S5c and d, respectively.

## Acknowledgements

We are grateful to M. Tarama, T. Yamamoto and D. Sipp for critical reading of this manuscript, and all member of Laboratory for Physical Biology for discussion. This work was supported by JSPS KAKENHI Grant Numbers JP17J05667 (to M.H.); JP16K17777 and JP19K03764 (to T.H.); JP26610129 (to H.K.); JP19H00996 (to T.S.); JST CREST grant number JPMJCR1852, Japan (T.S.)

## Author contributions

M.H. and T.S. designed the research. M.H. and Y.W. performed the experiments, M.H. and T.S. analyzed the data, H.K. provided KI cell and TgrB1^-^ KI cell and supervised the experiments, T.H. developed and performed the numerical simulation, and M.H., T. H., H. K. and T. S. participated in writing the manuscript.

## Data availability

The data that support the findings of this study are available from the corresponding author upon reasonable request.

---

**Supplementary Figure 1.**
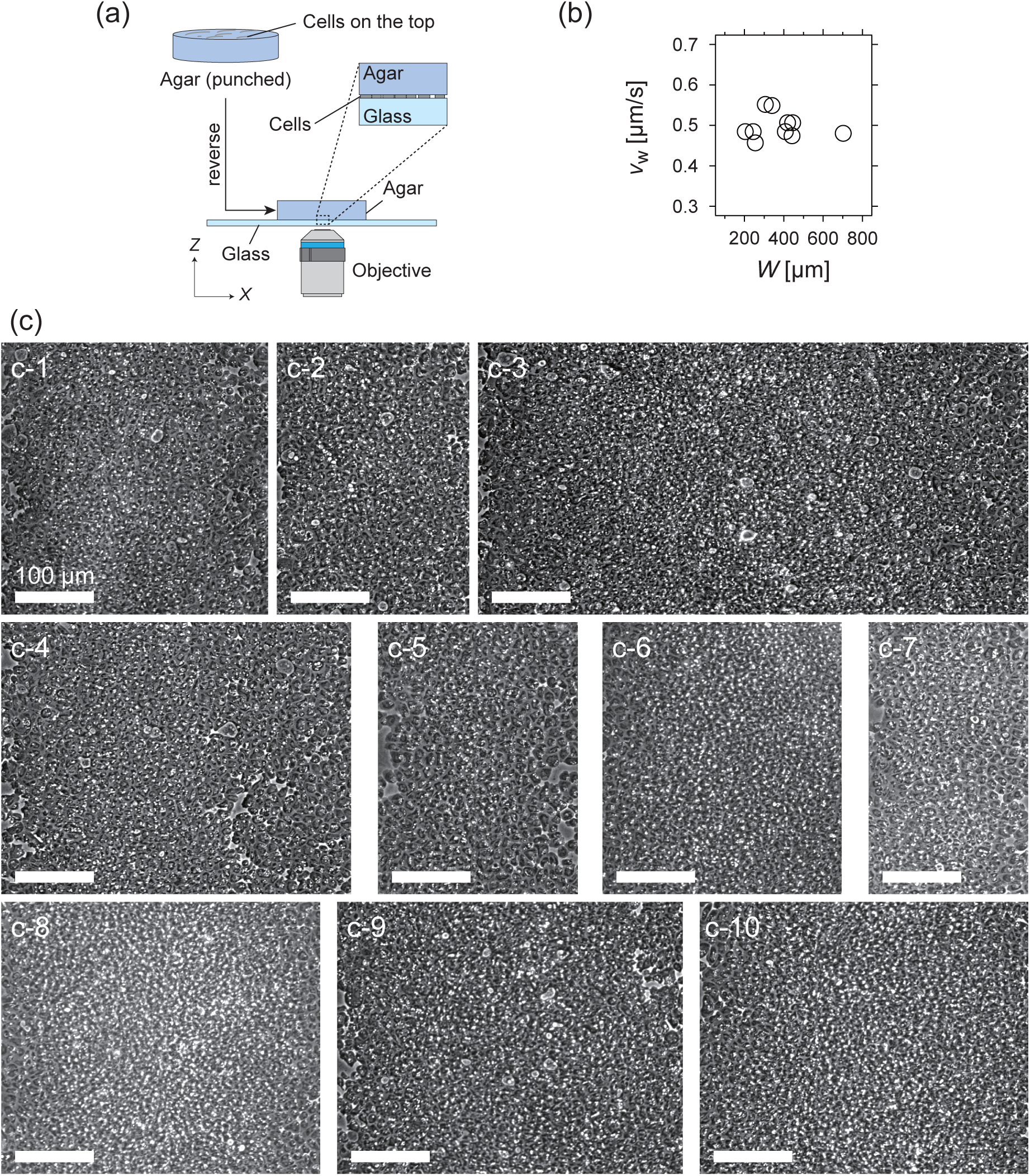
(a) Experimental setup for the microscopic observation of the polar pattern formation. (b) Relationship between the propagation speed *v*_w_ and size of the wave *W*. (c) Snapshots of propagating band. The number of waves studied is N =10. (c-1) *W* = 340 µm; (c-2) *W* = 244 µm; (c-3) *W* = 702 µm; (c-4) *W* = 445 µm; (c-5) *W* = 255 µm; (c-6) *W* = 306 µm; (c-7) *W* = 204 µm; (c-8) *W* = 408 µm; (c-9) *W* = 442 µm; (c-10) *W* = 419 µm.

**Supplementary Figure 2.**
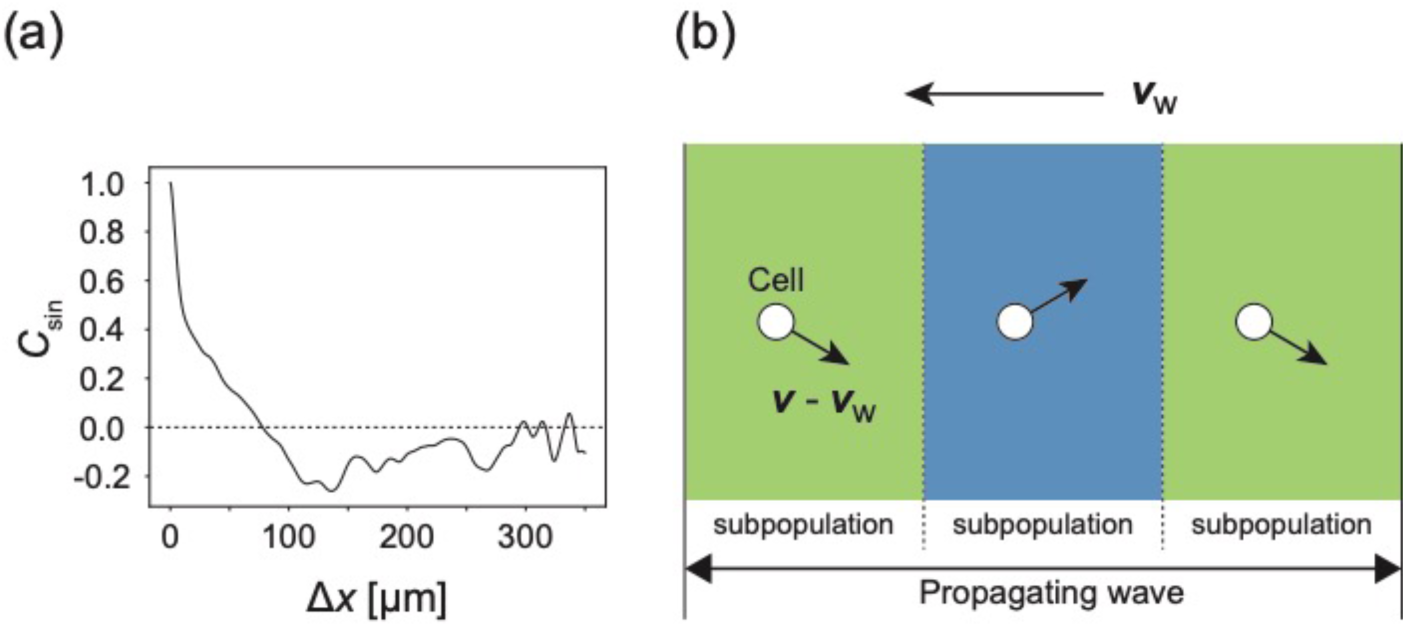
(a) **Autocorrelation function of transverse motion with respect to the wave propagation direction** *C*_sin_ (Δ*x*), indicating that the typical width of the stripe was around 125 µm. (b) Illustration of the cell migration within the formed stripes in the co-moving frame. Arrows from cells indicate the direction of motion relative to the velocity vector of wave propagation.

**Supplementary Figure 3.**
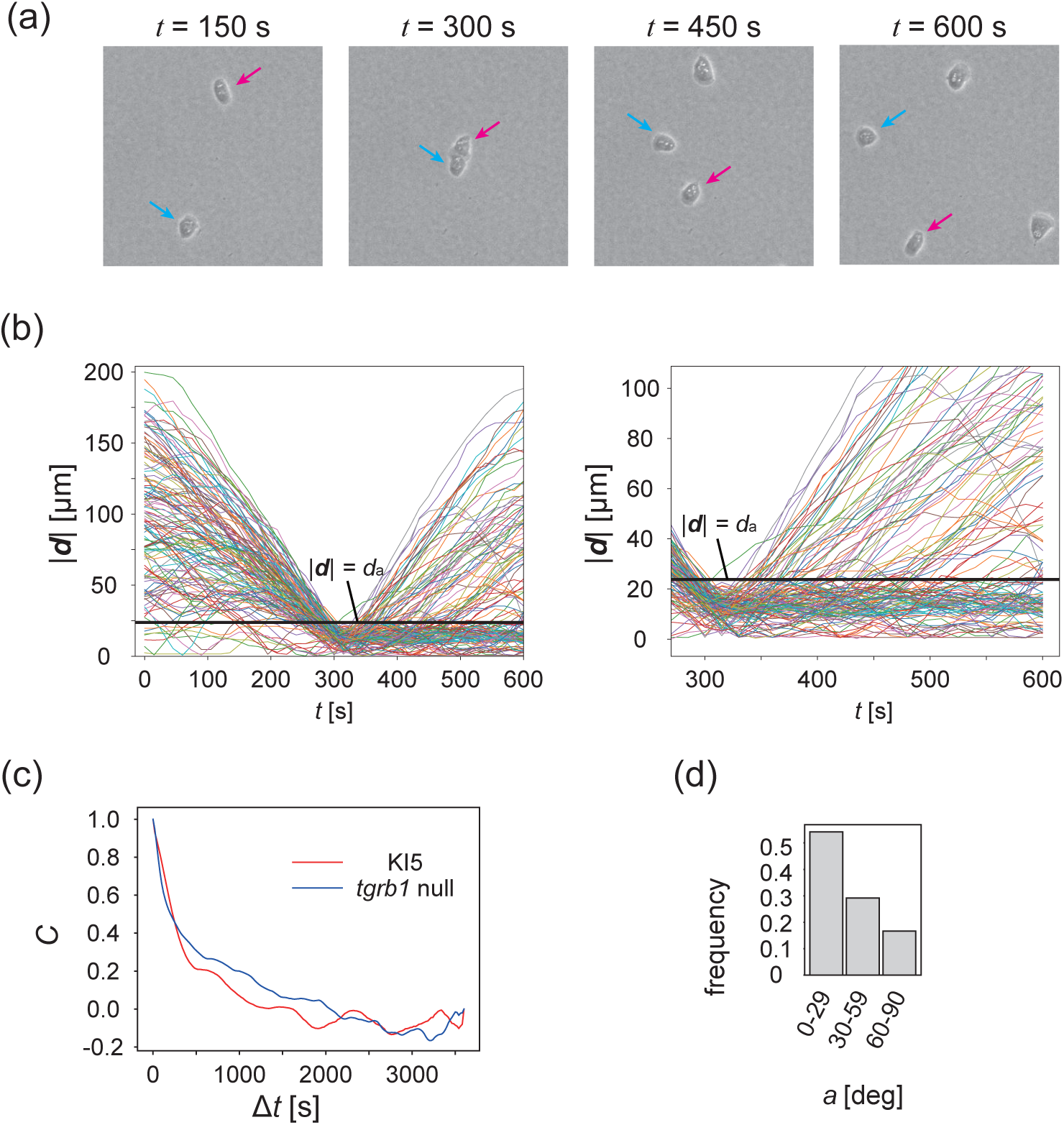
(a) Another example of time evolution of collision of two cells. (b) Time series of distance between two cells |***d***|. Each line is a data obtained from different cell pairs. Black horizontal line means |***d***| = *d*_a_. Time series of |***d***| after the collision (*t* = 300) is magnified in the right panel. (c) The velocity auto-correlation functions *C*(*τ*) of the isolated cells. Red line: KI cell. Blue line: *tgrb1* null mutant. Histogram of the CFL angle *a* for the *tgrb1* null mutant cells that contact each other for >300 sec.

**Supplementary Figure 4.**
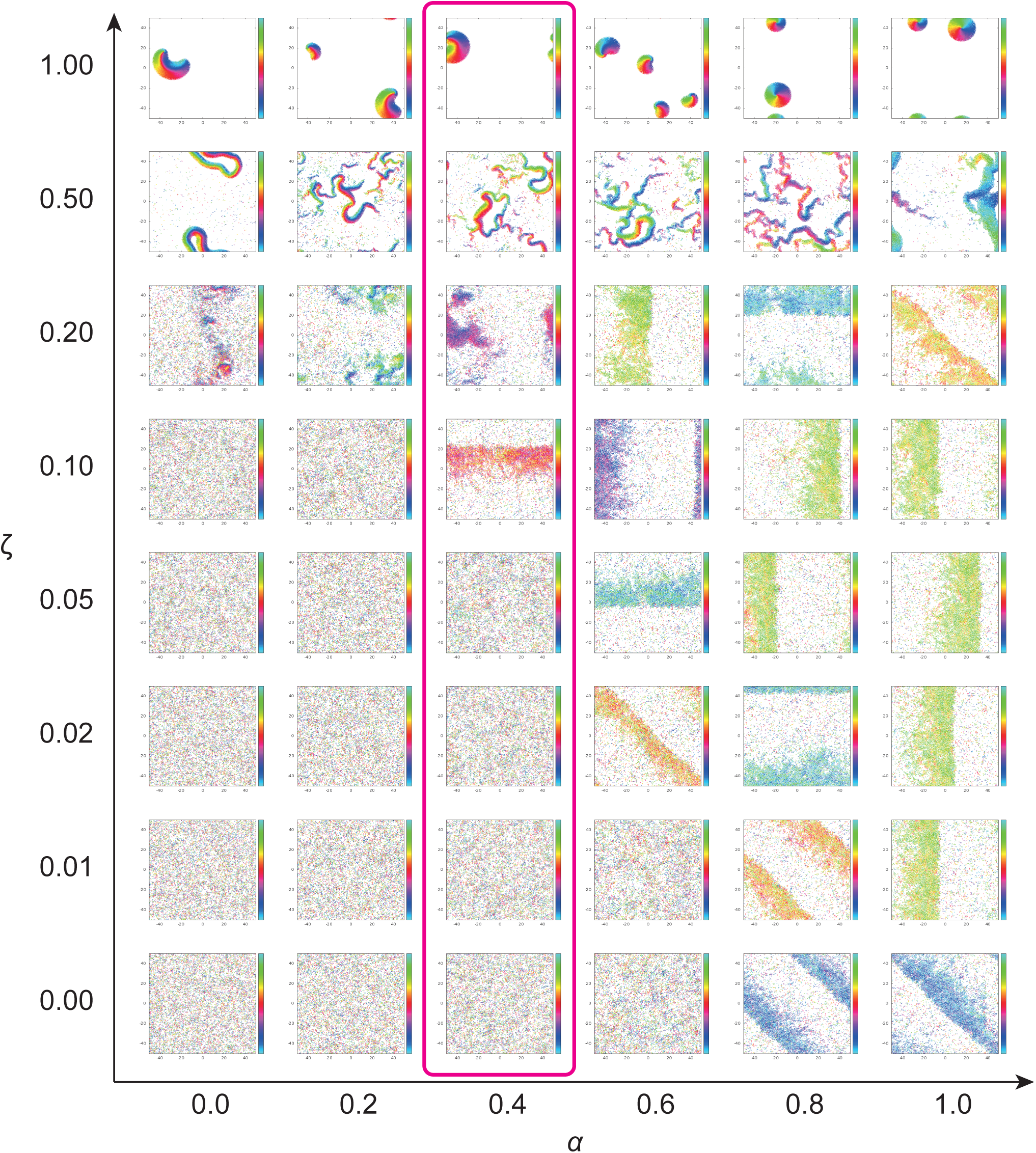
Phase diagram of the agent-based simulation for changes of CFL *α* and the alignment effect *ζ*. A region where α = 0.4 is surrounded by the pink line.

**Supplementary Figure 5.**
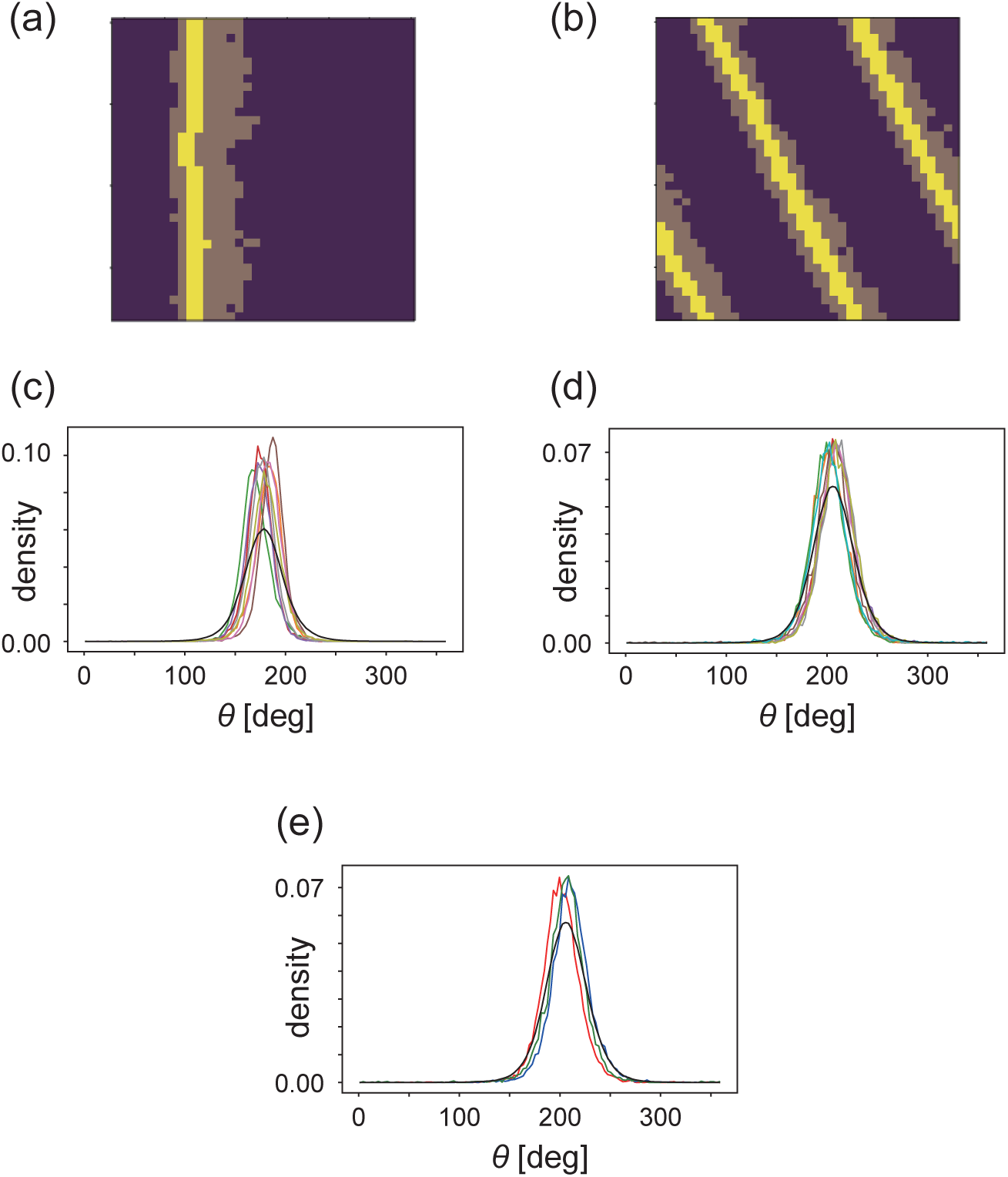
(a, b) The regions of interest used in the analyses (bright yellow) within the wave (dark yellow) for (a) CFL-induced and (b) alignment-induced waves, respectively. Analysis method to pick up these ROIs is found in Methods. (c, d) Pdfs of the migration direction within the wave region with the top eight and nine peak probability densities for (c) CFL-induced and (d) alignment-induced waves, respectively. (e) Typical pdfs for alignment-induced waves extracted from (d).

**Supplementary Movie 1** | Macroscopic observation of the propagating bands.

The movie was taken every 15 min for 28.5 hours. Video acceleration: 11400 × real time.

**Supplementary Movie 2** | Microscopic observation of the propagating bands.

The movie was taken every 15 s for 1.66 hours. Video acceleration: 230 × real time.

**Supplementary Movie 3** | Propagating band with overlaying the coloring based on the optical flow analysis.

The movie was taken every 3 s for 39.5 min. Video acceleration: 151 × real time.

**Supplementary Movie 4** | Migration of the KI cells in the low-density region.

The movie was taken every 15 s for 2 hours. Video acceleration: 378 × real time.

**Supplementary Movie 5** | A binary collision of the KI cells in the low-density assay.

The movie was cropped from the movie of the low-density assay with a length of 10.25 min. Video acceleration: 153 × real time.

**Supplementary Movie 6** | Macroscopic observation of the population of *tgrb1* null mutant.

The movie was taken every 15 min for 28.5 hours. Video acceleration: 11400 × real time.

**Supplementary Movie 7** | Microscopic observation of the population of *tgrb1* null mutant. The movie was taken every 15 s for 1.25 hours. Video acceleration: 225× real time.

**Supplementary Movie 8** | Propagating band formation generated in the agent-based simulation.

The color code indicates the migration direction of individual particle as shown in Fig 3**c**. Arrows indicate the direction of polarity.

## References

Ballerini M, Cabibbo N, Candelier R, Cavagna A, Cisbani E, Giardina I, Lecomte V, Orlandi A, Parisi G, Procaccini A, Viale M, Zdravkovic V. 2008. Interaction ruling animal collective behavior depends on topological rather than metric distance: Evidence from a field study. Proc Natl Acad Sci USA 105:1232–1237. doi:10.1073/pnas.0711437105

Butt T, Mufti T, Humayun A, Rosenthal PB, Khan S, Khan S, Molloy JE. 2010. Myosin Motors Drive Long Range Alignment of Actin Filaments. J Biol Chem 285:4964–4974. doi:10.1074/jbc.M109.044792

Chaté H, Ginelli F, Grégoire G, Raynaud F. 2008. Collective motion of self-propelled particles interacting without cohesion. Phys Rev E 77:046113. doi:10.1103/PhysRevE.77.046113

Dormann D, Parent CA. 2002. Visualizing PI3 kinase-mediated cell-cell signaling during Dictyostelium development. Curr Biol 12:1178–1188.

Fujimori T, Nakajima A, Shimada N, Sawai S. 2019. Tissue self-organization based on collective cell migration by contact activation of locomotion and chemotaxis. Proc Natl Acad Sci USA 116:4291–4296. doi:10.1073/pnas.1815063116

Ginelli F, Peruani F, Bär M, Chaté H. 2010. Large-Scale Collective Properties of Self-Propelled Rods. Phys Rev Lett 104:184502. doi:10.1103/PhysRevLett.104.184502

Haeger A, Wolf K, Zegers MM, Friedl P. 2015. Collective cell migration: guidance principles and hierarchies. Trends in Cell Biology 25:556–566. doi:10.1016/j.tcb.2015.06.003

Hashimura H, Morimoto YV, Yasui M, Ueda M. 2019. Collective cell migration of Dictyostelium without cAMP oscillations at multicellular stages. Commun Biol 2:34. doi:10.1038/s42003-018-0273-6

Hiraiwa T. 2019. Two types of exclusion interactions for self-propelled objects and collective motion induced by their combination. Phys Rev E 99:012614. doi:10.1103/PhysRevE.99.012614

Hiraiwa T, Nagamatsu A, Akuzawa N, Nishikawa M, Shibata T. 2014. Relevance of intracellular polarity to accuracy of eukaryotic chemotaxis. Phys Biol 11:056002. doi:10.1088/1478-3975/11/5/056002

Hirose S, Benabentos R, Ho H-I, Kuspa A, Shaulsky G. 2011. Self-recognition in social amoebae is mediated by allelic pairs of tiger genes. Science 333:467–470. doi:10.1126/science.1203903

Hirose S, Santhanam B, Katoh-Kurosawa M, Shaulsky G, Kuspa A. 2015. Allorecognition, via TgrB1 and TgrC1, mediates the transition from unicellularity to multicellularity in the social amoeba Dictyostelium discoideum. Development 142:3561–3570. doi:10.1242/dev.123281

Kida Y, Pan K, Kuwayama H. 2019. Some chemotactic mutants can be progress through development in chimeric populations. Differentiation 105:71–79. doi:10.1016/j.diff.2019.02.001

Kuwayama H. 2013. Biological soliton in multicellular movement. Sci Rep 3:2272. doi:10.1038/srep02272

Kuwayama H, Ishida S, Van Haastert PJ. 1993. Non-chemotactic Dictyostelium discoideum mutants with altered cGMP signal transduction. J Cell Biol 123:1453–1462. doi:10.1083/jcb.123.6.1453

Kuwayama H, Obara S, Morio T, Katoh M, Urushihara H, Tanaka Y. 2002. PCR-mediated generation of a gene disruption construct without the use of DNA ligase and plasmid vectors. Nucleic Acids Res 30:E2–2. doi:10.1093/nar/30.2.e2

Li B, Sun SX. 2014. Coherent motions in confluent cell monolayer sheets. Biophys J 107:1532–1541. doi:10.1016/j.bpj.2014.08.006

Li C-LF, Chen G, Webb AN, Shaulsky G. 2015. Altered N-glycosylation modulates TgrB1- and TgrC1-mediated development but not allorecognition in Dictyostelium. J Cell Sci 128:3990–3996. doi:10.1242/jcs.172882

Li D, Wang Y-L. 2018. Coordination of cell migration mediated by site-dependent cell– cell contact. Proc Natl Acad Sci USA 115:10678–10683. doi:10.1073/pnas.1807543115

Marchetti MC, Joanny J-F, Ramaswamy S, Liverpool TB, Prost J, Rao M, Simha RA. 2013. Hydrodynamics of soft active matter. Rev Mod Phys 85:1143–1189. doi:10.1103/RevModPhys.85.1143

Ohta T, Yamanaka S. 2014. Soliton-like behavior of traveling bands in self-propelled soft particles. Progress of Theoretical and Experimental Physics 2014:11J01–0. doi:10.1093/ptep/ptt111

Solon AP, Chaté H, Tailleur J. 2015. From Phase to Microphase Separation in Flocking Models: The Essential Role of Nonequilibrium Fluctuations. Phys Rev Lett 114:068101. doi:10.1103/PhysRevLett.114.068101

Sumino Y, Nagai KH, Yoshikawa K, Chaté H, Oiwa K. 2012. Large-scale vortex lattice emerging from collectively moving microtubules. Nature 483:448–452. doi:10.1038/nature10874

Suzuki R, Weber CA, Frey E, Bausch AR. 2015. Polar pattern formation in driven filament systems requires non-binary particle collisions. Nat Phys 11:839–843. doi:10.1038/nphys3423

Szabó B, Szöllösi GJ, Gönci B, Jurányi Z, Selmeczi D, Vicsek T. 2006. Phase transition in the collective migration of tissue cells: Experiment and model. Phys Rev E 74:061908. doi:10.1103/PhysRevE.74.061908

Takeuchi R, Tamura T, Nakayashiki T, Tanaka Y, Muto A, Wanner BL, Mori H. 2014. Colony-live--a high-throughput method for measuring microbial colony growth kinetics--reveals diverse growth effects of gene knockouts in Escherichia coli. BMC Microbiol 14:171. doi:10.1186/1471-2180-14-171

Vicsek T, Czirók A, Ben-Jacob E, Cohen I, Shochet O. 1995. Novel Type of Phase Transition in a System of Self-Driven Particles. Phys Rev Lett 75:1226–1229. doi:10.1103/PhysRevLett.75.1226

Vicsek T, Zafeiris A. 2012. Collective motion. Physics Reports 517:71–140. doi:10.1016/j.physrep.2012.03.004

Weber CA, Semmrich C. 2010. Polar patterns of driven filaments. Nature 467:73–77. doi:10.1038/nature09312

Williams JG. 2010. Dictyostelium finds new roles to model. Genetics 185:717–726. doi:10.1534/genetics.110.119297

Wioland H, Woodhouse FG, Kessler JO, Goldstein RE. 2013. Confinement stabilizes a bacterial suspension into a spiral vortex. Phys Rev Lett 110:268102. doi:10.1103/PhysRevLett.110.268102

Zhang HP, Be’er A, Florin E-L, Swinney HL. 2010. Collective motion and density fluctuations in bacterial colonies. Proc Natl Acad Sci USA 107:13626–13630. doi:10.1073/pnas.1001651107

